# Neodymium-Doped Nanocrystals for Diffraction-limited *in vitro* Temperature Sensing

**DOI:** 10.1101/2024.02.08.579538

**Authors:** M. Bravo, S. Yang, D. Wen, S. Krzyzowska, F. Taemaitree, S. Zaman, B. Fortuni, S. Rocha, P. Mulvaney, H. Uji-i, S. Brooke, J. Hutchison

## Abstract

Localized hyperthermia is a promising approach to cancer therapy. However, its clinical potential is limited by heterogeneous heat distribution within tumors, and advanced methods to measure temperature at the sub-micron level are therefore required. To address this challenge, luminescent nanothermometers, such as lanthanide-doped nanocrystals (Ln-NC) operating in the near-infrared (NIR), have been investigated for accurate spatiotemporal thermal monitoring. In this study, the synthesis of neodymium-doped, sodium yttrium fluoride nanocrystals (Nd-NCs) was optimized to achieve high photoluminescence (PL) intensity by adjusting the dopant concentration and by shelling with inert layers. Standard curves for luminescence-based temperature readout were developed using ratiometric analysis of the temperature-dependent PL spectra in the 850-920 nm biological window, showing excellent linearity and high thermal sensitivity. A silica shell was added to the particles and shown to confer excellent aqueous stability and biocompatibility in A549 lung cancer cells. Finally, luminescent thermal readout was demonstrated *in vitro* in A549 cells by spectrally resolving the diffraction-limited luminescence spots at a single-particle scale over a clinically relevant temperature range from 20-50 °C. The application of the developed nanothermometer as preclinical tools for NP-HT characterization could provide crucial information on the therapeutic temperature achieved in and around the tumor area. This could be key to optimizing NP properties and therapeutic parameters, for the development of viable hyperthermal cancer treatments.

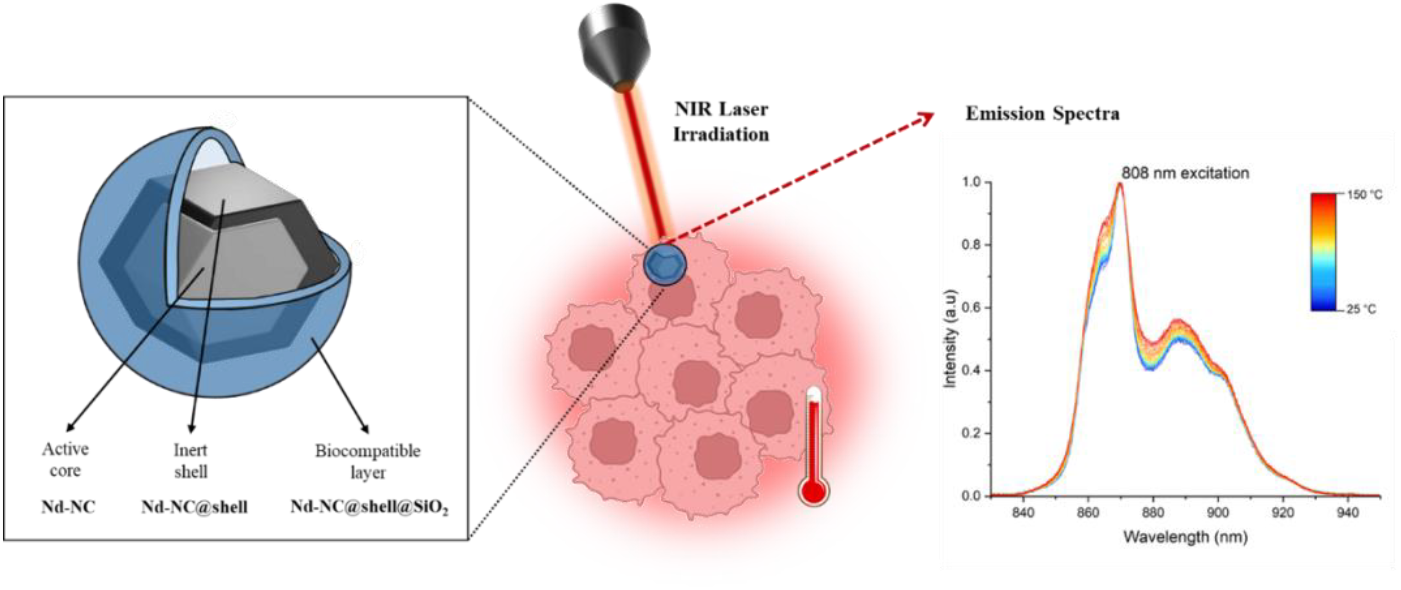

## 1. Introduction

Nanoparticle hyperthermia (NP-HT) is an emerging cancer therapy that uses heat-generating NPs to locally increase temperature to induce a heat shock reaction in cancer cells^1,2^. NPs within the 20 to 200 nm range can extravasate leaky tumor blood vessels and accumulate at the tumor site via the enhanced permeability and retention (EPR) effect^3^. Surface-functionalization of NPs with moieties that specifically bind to tumor biomarkers further aids active tumor targeting^7,8^. This tumor-specific accumulation increases therapeutic specificity and reduces off-target toxicity to healthy tissues^9^. While NP-HT provides a promising alternative to conventional approaches, several challenges remain in translating promising results to the clinic. One of the main hurdles is the lack of accurate temperature readouts during treatment, leading to heterogeneous heat distribution within and around the tumor area, and limited control over therapeutic outcomes^3^. When temperatures exceed 45 °C, tumor destruction is achieved via thermal ablation as a result of irreversible coagulation and protein denaturation. Therefore, thermal ablation can be explored as a standalone therapy^9,10^. Lower temperatures (40-45 °C), induce reversible physiological changes, including enhanced blood profusion and vascular permeability at the tumor site, to higher drug delivery efficiency, and tumor oxygenation. In this case, HT can be explored as an adjuvant strategy, due to enhanced tumor sensitivity to other therapies^11,12^. Considering the thermal effect observed for HT, an accurate thermal control within the tumor mass is crucial, to optimize anti-tumor action, while keeping the surrounding tissue within biologically safe levels^3^. To address this, accurate spatiotemporal and thermal profiling in solid tumors during HT is essential for designing effective therapeutic outcomes with minimal side effects, by enabling the fine-tuning of therapeutic parameters^9,13^.

Most HT procedures to date employ contact thermometers, such as thermocouples and fiber optic sensors, which offer high thermal resolution but are invasive and limited to single-point thermal readouts within a tumor^14–16^. Infrared (IR) imaging is a non-contact alternative with high thermal and spatial resolution^16^, but is restricted to superficial temperature sensing due to tissue emissivity^17,18^. Once again nanoparticles may provide a promising solution, enabling non-contact, real-time temperature readouts even from sub-cutaneous tumors during HT^19,20^. Luminescent nanothermometers display relevant temperature-dependent emission in the physiological and above-physiological temperature range, which can be used to derive local temperature with sub-micron spatial resolution^16^. Over the past couple of decades, different temperature-sensitive nanoprobes have been developed. In particular, lanthanide-doped nanocrystals (Ln-NCs) have emerged as ideal candidates, since they exhibit low phototoxicity, no photobleaching, high signal-to-noise ratio, and ease of functionalization for improved biocompatibility^21^. Ln-NCs comprise an optically transparent crystal host, that is doped with varying concentrations of one or more lanthanide ions (Ln^3+^) ^22^. Neodymium (Nd^3+^) is one of the most widely studied Ln^3+^ for bio-sensing and bio-imaging applications, mainly because it exhibits excitation and emission lines within the first biological transparency window in the NIR (800-950 nm)^16^. In this spectral region, light absorption and scattering by biological tissue is reduced, improving the penetration of optical excitation and luminescent temperature readout^16,19^.

In this work we explore the design of biocompatible Nd^3+^-doped NCs (Nd-NCs) for precise temperature monitoring inside biological environments. Sodium yttrium fluoride (NaYF_4_) was chosen as the host material, due to its high optical transparency, and its low phonon energies which minimize nonradiative multiphonon relaxation processes that quench dopant luminescence^23,24^. Additionally, NaYF_4_ offers a robust matrix that can accommodate the lanthanide ions without significantly disrupting the lattice structure^25^. Nd-NCs were synthesized via a thermal decomposition method and their photoluminescence (PL) was optimized by tuning the dopant concentration and adding a protective inert NaYF_4_ shell (Nd-NC@shell). To reduce cytotoxicity and ensure aqueous stability, the as-synthesized Nd-NC@shell particles were encapsulated inside a biocompatible silica (SiO_2_) layer (Nd-NC@shell@SiO_2_). NIR excitation (785 nm and 808 nm lasers) was used to generate temperature-dependent, multi-peaked Nd emission between 830-950 nm, from which we can construct standard curves of temperature vs luminescence peak ratio. Nd-NC@shell@SiO_2_ particles were efficiently incorporated into A549 cancer cells without compromising cell viability. Most importantly, luminescence imaging and microspectroscopy at low particle loadings were used to resolve the diffraction-limited emission point spread functions from single particles within A549 cells. Single particle spectra are found to vary with temperature over the range 20-50 °C *in vitro*, demonstrating the applicability of our NPs as local temperature probes in biological environments. This work provides crucial information on the use of Ln-doped nanothermometers in preclinical HT studies, and will be an essential tool for the rational design of next-generation nanomaterials for targeted light-mediated cancer therapies.

## 2. Materials and Methods

### 2.1. Materials

Sodium trifluoroacetate (Na(CF_3_COO)_3_, 98%), trifluoroacetic acid (TFA, 99%), Yttrium(III) oxide (nanopowder, <50 nm particle size), 1-Octadecene- (ODE, 90%), oleic acid (OA, 90%), Tetraethyl Orthosilicate (TEOS, 98%), IGEPAL® CA-630, Ammonium hydroxide (NH_4_OH, 28-30%), Cyclohexane (≥ 99.8%), and Poly(methyl methacrylate) (PMMA), were purchased from Sigma-Aldrich; Neodymium oxide was purchased from Koch-Light Laboratories Ltd; Polyethylene Glycol (PEG, MW 8000) was purchased from MP Biomedicals; Dulbecco’s modified eagle medium (DMEM), Gentamicin, Dulbecco’s phosphate Buffered saline (PBS, no calcium, no magnesium), Formaldehyde (4% in PBS), trypsin-EDTA (0.5%), and Hank’s balanced salt solution (HBSS, no phenol red) were purchased from Wako Pure Chemical Industries; Triton X-100 (0.1%) was purchased from Nacalai Tesque, INC; Alexa Fluor 488 Palloidin 488, Vybrant® DiO cell-labeling solution, and L-glutamine were purchased from Thermofisher Scientific; Fetal bovine serum (FBS) was purchased from Gibco; Cell Counting Kit-8 (CCK-8) was purchased from Dojindo; Milli-Q water is from Millipore. All the chemicals were used without further purification.

### 2.2. Methods

#### Synthesis of Core and Core-shell Nanocrystals

Lanthanide fluoroacetate precursors were prepared by dissolving 500 mg of Nd_2_O_3_ or Y_2_O_3_ in a 1:1 mixture of Milli-Q water and trifluoroacetic acid (5 mL each). The solution was heated to 80°C and stirred (500 rpm) until transparent. Residual solvents were evaporated at 120°C, yielding Nd(CF_3_COO)_3_ and Y(CF_3_COO)_3_ powders. Lanthanide-doped nanocrystals were synthesized via thermal decomposition of the trifluoroacetate precursors^26^. Briefly, x% mmol Nd(CF_3_COO)_3_ and (1-x)% 1.25 mmol Y(CF_3_COO)_3_ (total 1.25 mmol, x = 1, 2, 5 or 10%) were mixed with Na(CF_3_COO)_3_ (1.25 mmol) in a 3-necked 100 mL round flask and dispersed in 1-octadecene and oleic acid (10 mL each). The mixture was heated to 100°C while stirring, degassed, and allowed to react for 1.5 hours at 300°C under N_2_. After synthesis, the mixture was cooled to RT, and NPs were precipitated by a mixture of 4:1 acetone and hexane, and collected by centrifugation (3,000 rcf, 10 min). The NPs were then washed twice in 1:10 hexane and ethanol (3,000 rcf, 10 min), and dispersed in toluene (3 mL), yielding a similar NP concentration for every dopant concentration, of around (42 ± 3) mg/mL. For core-shell NP preparation, Y(CF_3_COO)_3_ and Na(CF_3_COO)_3_ (1 mmol) were dissolved in octadecene and oleic acid (5 mL each). After core formation, the reaction temperature was lowered to 250°C, and the shell precursor solution was added dropwise to the flask under stirring. The temperature was then raised to 305°C and allowed to react until transparent (∼ 1 hour). The mixture was cooled to RT, and the prepared NaYF_4_:Nd@NaYF_4_ NPs (hereafter referred to as Nd-NC@shell) were washed and precipitated as previously described, yielding a stable NP solution in toluene of (65 ± 5) mg/mL^27^. Detailed information about the synthesized NPs can be found in Table S1.

##### Silica-coated Nanocrystals

Nd-NC@shell NPs were coated with silica (SiO_2_) using a modified water-in-oil microemulsion protocol^28^. Briefly, IGEPAL® CA-630 (1.3 ml) was dispersed in cyclohexane (10 ml) by ultrasonication. 1 mL of diluted Nd-NC@shell solution (320 uL NCs in 670 uL cyclohexane) (∼ 20 mg/mL) was added to the mixture and stirred for 30 min. Then, ammonia (30%, 150 µL) was added dropwise, and the solution was stirred (500 rpm, 5 min) until clear. Lastly, 320 µL of TEOS solution (1:4 in cyclohexane) was added dropwise, homogenized (500 rpm, 5 min) and left undisturbed overnight to encourage hydrolysis and condensation of the silica precursor. The resulting Nd-NCs@shell@SiO_2_ were precipitated in 10 mL ethanol (1,000 rcf, 20 min) and isolated via centrifugation in ethanol 3 times, followed by 2 washes in Milli-Q water (11,000 rpm, 10 min). Nd-NC@shell@SiO_2_ were then resuspended in 320 µL of Milli-Q water.

##### NP structure/composition/charge characterization

As-synthesized colloidal dispersions were diluted in a 1:200 ratio and 6 μL of solution was cast onto TEM grids (300 mesh copper grid) and dried at room temperature. TEM images were collected on an FEI TecnaiF20 microscope (200 kV) (Ian Holmes Imaging Center, Bio21) equipped with an energy dispersive X-ray (EDX) detector for sample characterization, including size and chemical composition analysis. Crystal structure analysis was performed by comparing diffraction peaks of the synthesized NPs with X-Ray Diffraction patterns (XRD) of standard host lattices. The plots were obtained by determining the Bragg angle (θ) and plotting the intensity of the diffraction peaks against 2θ. To do so, the Bragg’s Law was used to determine the value of θ, where n equals 1, λ is the wavelength of the X-ray and d is the distance between the lattice plane (Equation 1)^31^.

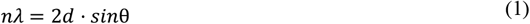

NP charge was characterized by zeta potential measurements (Zetasizer, Malvern). The morphology of NPs was further characterized using SEM (HITACHI HD-2000).

##### Optical Characterization/ Emission Spectra Collection

NIR photoluminescence measurements were performed using a homebuilt system. An 808 nm diode laser (SDL-808-LM-100T, Shanghai Dream Lasers Technology Co., Ltd) and a 785 nm diode laser (SDL-785-LM-100T, Shanghai Dream Lasers Technology Co., Ltd) were used to excite Ln-doped nanocrystals via an inverted optical microscope (Olympus IX-71) in epifluorescence configuration using a half-silvered mirror. The excitation power was around 0.15 mW. The NP luminescence was collected by a 20x objective (NA = 0.45), separated from excitation light with longpass filters, then guided into a spectrograph/silicon CCD couple (Acton 2150/Princeton 1024B). The luminescence was recorded in the range 725-1000 nm. For generating standard curves for luminescence peak ratio vs temperature, 15 µL of sample was drop-cast onto a round coverslip and placed inside a temperature-controlled stage (Linkam THMS600). Emission spectra were recorded over a temperature range from 25°C to 150°C, at 5°C intervals. Within the therapeutically relevant range (35-60°C), data was collected at 1°C temperature steps. Acquisition time varied between samples, and PL was standardized using calibration curves obtained for Intensity vs . Acquisition time plots (Figure S1). Spectra were analyzed using Origin 2024b, and intensity data displayed as means ± standard deviations with error bars indicating one standard deviation.

##### Cell Culture

A549 cells were cultured in 25 cm^2^ culture flasks at 37° C under 5% CO_2_ atmosphere. Cells were maintained in DMEM medium with 10% FBS, 1% L-glutamax, and 0.1% gentamicin. For fluorescence microscopy experiments, the cells were seeded in 31-mm, glass-bottom dishes (Cellvis, Mountain View, CA, USA) and grown until ∼80% confluency before adding the NPs.

##### Cytotoxicity Studies

NP toxicity was assessed using the cell counting kit-8 (cck-8) assay kit. A549 cells were seeded in a 96-well plate at a density of 5×10^4^ cells/well. The next day, 200 µL of increasing concentrations of Nd-NPs@mSi (1, 5, 10, 15, 25 and 50µg/mL) were added to the cells. Three biological replicates were prepared for each condition. Cells incubated with nanoparticles were washed with PBS after 6 h of nanoparticle incubation to remove the non-internalized NPs. After washing, fresh DMEM was added and the sample was incubated for 24 additional hours. After incubation, the cells were washed with medium and 100 µL CCK-8 solution was added to each well and incubated for 1 hour. Absorption values were measured with a microplate reader and normalized to the OD450−OD620 value for the untreated cells.

##### Single NP emission

*In vitro* single NP luminescence studies were performed using a microspectroscopy apparatus consisting of 808 nm diode lasers coupled to an inverted microscope in epi-fluorescence configuration. The excitation light was focused at the back focal plane of the objective to give wide-field excitation. Emission was collected through the same objective, separated from excitation light by long-pass filters, and imaged on a silicon EMCCD camera. Emission from specific diffraction-limited spots was isolated by a slit and spectrally resolved. For *in vitro* studies, A549 lung cancer cells were incubated with Nd-NC@shell@SiO_2_ (50 µg/mL). Cells were stained with Alexa Fluor 488 Phalloidin. In short, cells were seeded in 31-mm glass-bottom dishes and grown overnight. Next, the cells were fixed with paraformaldehyde (4%), and the membrane was permeabilized with Triton X-100 (0.1%), for 15 min. The sample was carefully washed with PBS (1×) between each step. After washing, 200 µL of phalloidin 488 in 3% BSA solution (1:500 dilution) was added to the dish and the sample was incubated at RT for 30 min. After incubation, the samples were washed twice with PBS (1×).

##### Statistical Analysis

The data are displayed as means ± standard deviations and error bars indicate standard deviation. A randomization test was used to compare any two groups of values and performed using T-TEST function in Microsoft Excel. Statistical significance was reported as * p < 0.05, ** p < 0.01, and *** p < 0.001.

## 3. Results and discussion

### 3.1. Characterization of Nd-NC Nanoparticles

NaYF_4_:Nd, (Nd-NC) core NPs were synthesized via thermal decomposition of lanthanide precursors^26^. Different Nd^3+^ ion dopant percentages (1%, 2%, 5%, and 10%, with respect to the Y^3+^ ion concentration) were tested to investigate the impact of Nd^3+^ concentration on PL. The brightest cores were then shelled with an inert NaYF_4_ shell. The samples were stable in toluene for several months. Transmission electron microscopy (TEM) was used to determine the size and morphology of Nd-NC when 0, 1, 2, 5 or 10% Nd^3+^ ion dopant was used (Fig. 1a, and Fig. S2 a-d). The NPs displayed homogeneous size distribution around 30 nm, and polyhedral shape for all the different Nd^3+^ concentrations. The size distribution plot showed no significant size changes between samples (Fig. 1c). This is explained by the close match between the ionic radius of Nd^3+^ (*r*_Nd_^3+^=1.123 Å) and Y^3+^ (*r*_Y_^3+^=1.040 Å). The replacement of Y^3+^ ions by Nd^3+^ does not significantly impact the crystal structure^32,33^. However, the addition of the NaYF_4_ shell led to a significant size increase (∼25%) of 9 nm (shell thickness = 4.5 nm) (Fig. 1b, c). Elemental analysis was performed to assess the presence of Nd^3+^ and estimate its proportion via Energy Dispersive Spectroscopy (EDS) analysis (Fig. S2e). The obtained atomic percentages showed a strong (linear) correlation between the increasing amount of Nd^3+^ employed in the synthesis, and the final percentage of lanthanide ion in the host matrix (Fig. S2f), indicating successful doping.

**Figure 1.**
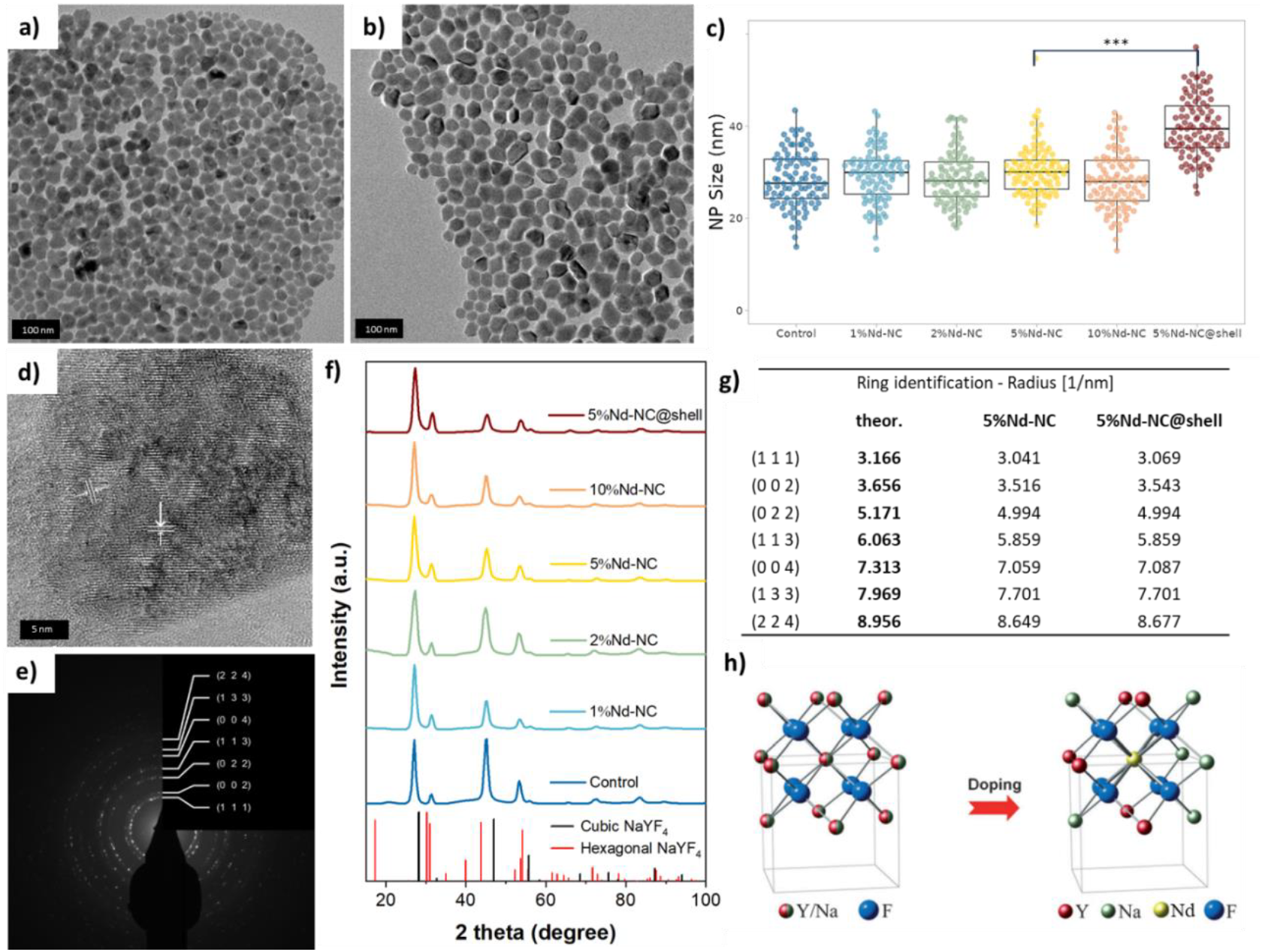
Transmission Electron Microscopy (TEM) images of (a) 5%Nd-NCs and (b) 5%Nd-NCs@shell. Scale bar = 100 nm; (c) Size distribution of Nd-NCs with different dopant concentrations and Nd-NC@shell. No significant changes in size were observed for different dopant concentrations (p > 0.05), but the size significantly increased with the addition of a NaYF_4_ shell (p < 0.001). NP size was estimated from 100 NPs; (d) High-resolution TEM image of a single 5%Nd-NC showing lattice fringes; (e) SAED of 5%Nd-NC showing the diffraction pattern for different planes (Miller indices noted) – SAED data was analyzed with crystbox^38^; (f) Diffraction pattern plot of the synthesized Nd-NPs (0, 1, 2, 5 and 10%), Nd-NC@shell, as well as the standard cubic and hexagonal NaYF_4_ standard host lattices, showing a close correlation between the diffraction peaks of synthesized NCs and the standard α-cubic host lattice (in black); (g) Ring identification for various crystal planes (Miller index noted) of standard cubic NaYF_4_, 5%Nd-NC, and 5%Nd-NC@shell, showing very close agreement, (h) Graphical representation of the structure of cubic NaYF_4_ and Nd-NCs (Adapted from ^39^).

High-resolution TEM images of individual Nd-NCs (Fig. 1d) revealed clear lattice fringes (3.5 Å spacing), indicating a highly crystalline structure in accordance with literature data on NaYF_4_ ^34,35^. NaYF_4_ host lattices can present two thermodynamically stable phases, cubic (isotropic) α-phase, and hexagonal (anisotropic) β-phase^36^. Selected area electron diffraction (SAED, Fig. 1e), was used to identify which of the phases correspond to the synthesized NCs. To do so, d-spacing values for the different NCs were converted to Bragg angle (θ) (Eq. 1, methods section). The intensity of the diffraction peaks was plotted against 2θ, enabling a comparison with X-ray diffraction (XRD) patterns of the standard cubic (COD-1517676) and hexagonal (COD-1517672) phases. The resulting plot shows a high overlap between the standard peak pattern of the α-cubic phase of NaYF_4_ and all the as-synthesized NCs (negative control, 1%, 2%, 5%, 10%) (Fig. 1f). Furthermore, the diffraction pattern plots of the synthesized NCs show narrow, intense peaks with no anomalous peaks, reflecting a well-defined crystalline structure with no impurities^31,36^. A small shift was observed for all the peaks compared to the standard NaYF_4_ phases, which could be attributed to lattice strain in the NPs compared to the bulk material^37^. D-spacing values were obtained for the different diffraction rings of the designed Nd-NCs, and were again in excellent agreement with the standards (Fig. 1g). All of this corroborating data confirms that the synthesized NCs NaYF_4_ matrix is in the α-cubic phase, and virtually unaffected by Nd^3+^ doping up to 10%.

### 3.2. Photoluminescent Properties

Luminescence nanothermometry in cellular environments benefits from brightly luminescent NPs that allow for effective emission readout from lower concentrations, reducing the risk of concentration-induced toxicity^40^. To optimize photoluminescence (PL), we studied the impact of Nd^3+^ concentration on the emission of the Nd-NCs. Emission spectra were collected for colloidal dispersions at room temperature, using two distinct excitation wavelengths within the first biological window (NIR-I BW): 785 nm and 808 nm, with constant laser power (0.15 mW). Nd^3+^ has a complex 4f^3^ electronic structure with 364 individual electronic states, which can be grouped into 41 multiplets of electronic energy levels^41. 4^F_3/2_ →^4^I_13/2_ (1350 nm) ^4^F_3/2_→^4^I_9/2_ (880 nm), ^4^F_5/2,_ ^2^H(2)_9/2_→^4^I_9/2_ (800nm), ^4^F_7/2,_ ^4^S_3/2_ →^4^I_9/2_ (740 nm) are amongst the most important excited multiplets (Fig. 2a). The first transition falls within the second BW, where light absorption and scattering by the biological tissue are even lower than for NIR-I BW^3^. However, detection efficiency at these wavelengths is poor, limiting applications^41^. Therefore, our study focused on emission in the NIR-I BW (830-950nm). In this range, excitation with 785/808 nm lasers, promotes electronic transitions from the ^4^I_9/2_ ground state multiplet to one of the excited states, followed by non-radiative relaxation to the metastable ^4^F_3/2_ state before emission occurs^41,42^.

**Figure 2.**
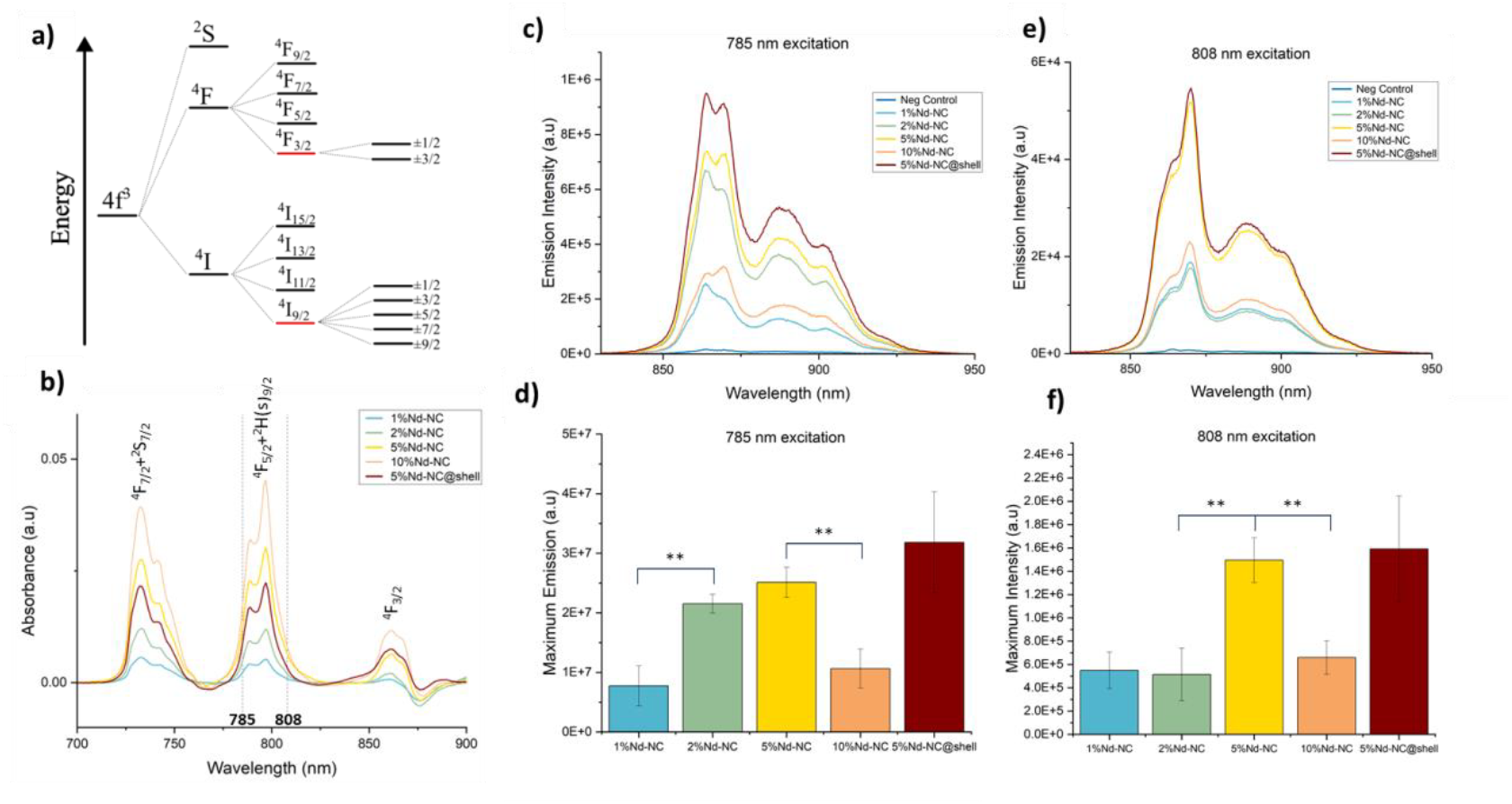
(a) Schematic energy level diagram showing the Stark-sublevels of ^4^F_3/2_; ^4^I_9/_2 involved in optical absorption and emission of Nd^3+^ after NIR excitation (Adapted from ^41^); (b) Absorbance spectra of Nd-NCs (1%, 2%, 5%, 10%) and Nd-NC@shell NPs. Absorbance increases with the dopant concentration and decreases with the addition of a shell (5%Nd-NC vs 5%Nd-NC@shell). All the samples show higher absorbance at 785 nm compared to 808 nm (p < 0.05); Emission spectra of all the NCs upon (c) 785 nm and (e) 808 nm excitation, respectively. Each spectrum is an average of 3 distinct samples (N=3); Bar graphs comparing the integrated area below the emission spectra for each sample upon (d) 785 nm and (f) 808 nm excitation. All the analyzed samples displayed similar concentrations ((42 ± 3) mg/mL, in toluene), therefore for both excitation wavelengths, 5% Nd exhibits the brightest per-particle PL (p < 0.05). The addition of an optically inert shell further improved emission intensity by 10-20%. Error bars indicate ± SD.

The absorption spectra for all samples showed overlapping peaks characteristic of the Nd^3+^ ion, correlated with the abovementioned transitions ^4^I_9/2_ →^4^F_7/2_+^2^S_7/2, 4_I_9/2_ →^4^F_5/2_+^2^H_9/2_ and ^4^I_9/2_ →^4^F_3/2_ transitions (Fig. 2b)^16,41,43^. The spectra revealed a concentration-dependent increase in absorption, with 10%Nd-NC showing the highest absorption at both 785 and 808 nm, with the addition of an inert shell leading to a slight decrease in optical absorption. Emission spectra analysis revealed four different downshifted peaks (Fig. S3a), resulting from transitions from different Stark sublevels of the ^4^F_3/2_ excited state to ^4^I_9/2_^16^. The PL spectra and integrated emission (830 to 950 nm) for the core Nd-NCs and the Nd-NC@shell NPs are compiled in Figure 2 (c-f). Differences in emission response were observed between the two excitation wavelengths. Nd-NCs displayed higher fluorescence under 785 nm excitation compared to 808 nm excitation (p < 0.05 for all samples). This correlation can be attributed to the higher absorption at 785 nm for all core and core-shell NPs (Fig. 2b), as all the core and core-shell NPs show higher absorption at 785 nm than 808 nm.

After 785 nm laser excitation (Fig. 2c, d), PL intensity increased with Nd^3+^ concentration up to 5% Nd^3+^, but decreased significantly (by 58%) for 10%Nd-NC, consistent with other literature reports^44,45^. In theory, an increasing dopant concentration should increase PL intensity due to an increase in the number of emitting centers. However, it has been shown that beyond a certain concentration threshold, the shorter distance between Nd^3+^ ions in the NaYF_4_ lattice promotes non-radiative cross-relaxation, which decreases overall PL^42,44,46^. A similar pattern was observed for 808 nm excitation (Fig. 2e, f). 5%Nd-NC displayed the highest emission intensity, followed by a 56% intensity loss for 10%Nd-NC. The negative control revealed no luminescence, confirming that the Nd^3+^ dopant is the only emitter in this spectral range. Based on these results, 5%Nd-NC was chosen as the optimal candidate for nanothermometry in biomedical applications, since its higher per-particle emission allows for the best emission readouts with the lowest NP concentrations. An inert (undoped) NaYF_4_ shell was added to the 5%Nd-NCs. As expected, PL was increased by 21% and 6% for 785 and 808 nm excitation, respectively. This enhancement is a result of reduced quenching caused by resonance energy transfer between Nd^3+^ optical transitions and C-H and O-H vibrational modes of the solvent, as well as reduction of surface defects near the active core^44,47^. The latter is especially prominent in aqueous environments as the O-H stretching mode of water is a powerful quencher of Ln^3+^ excited states^44,47^.

#### Temperature Dependence

To analyze the thermosensitive properties of the nanothermometers, PL emission spectra were collected at different temperatures (25-150°C) at 785 nm and 808 nm excitation (Fig. 3a, b), and data was baselined, and normalized ([0,1]) to the emission maximum in this region (Fig. S4a). Within the biologically and therapeutically relevant thermal window (35-60°C), data points were collected at 1°C intervals, with the remaining spectra collected at 5°C temperature increases. Nd-NC@shell showed clear temperature-dependent emission, as indicated by the color gradients at both wavelengths (Fig. 3a, d).

**Figure 3.**
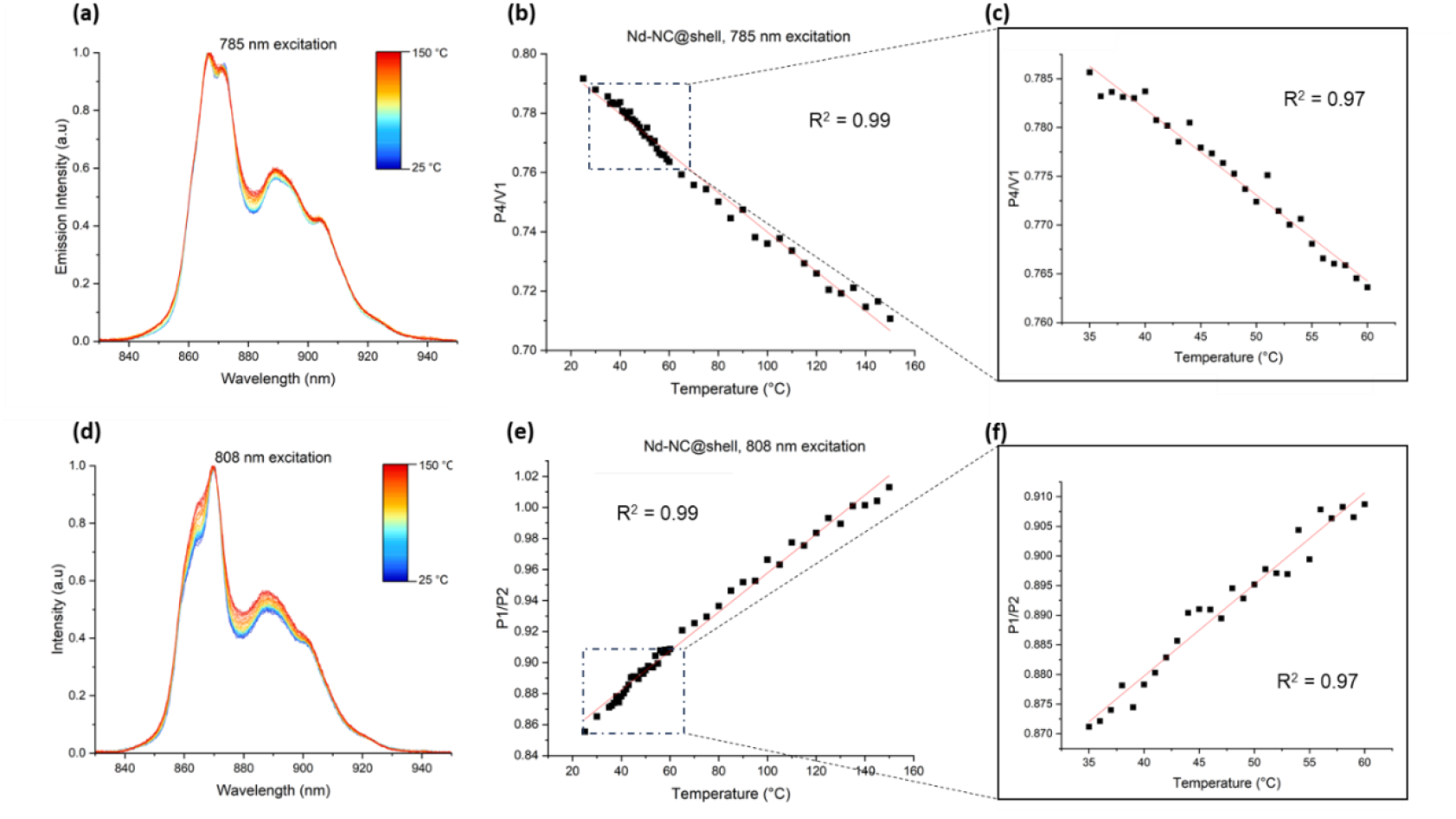
(a) Temperature dependence plot obtained for Nd-NC@shell NPs upon 785 nm and laser excitation. Emission spectra for all the samples were collected at increasing temperatures, starting at 25°C with a 5°C increase until 150°C. For the biologically relevant temperature range (35-60°C), spectra were collected at 1°C intervals. Data is shown as an average of 3 acquired spectra (N=3); (b) Calibration curves derived by calculating LIR (integrated wavelengths across 5 nm intervals of P4/V1), resulting in R^2^=0.99; (c) Zoomed in calibration curve focused on a HT-relevant temperature range (20-60°C) upon 785 nm excitation, also revealed high correlation (R^2^=0.97). (d) Temperature dependence plot obtained after 808 nm laser excitation; (e) Calibration curve derived from LIR (across 2 nm integrated wavelengths of P1/P2), showing high linear correlation (R^2^=0.99). (f) Zoomed in calibration curve within a biological and therapeutically relevant window at 808 nm excitation (R^2^=0.97).

Thermal calibration curves were constructed using luminescent intensity ratios (LIRs) of thermally coupled electronic states, a robust strategy for luminescence nanothermometry. This approach is independent of nanothermometer concentration, excitation intensity, and photobleaching^16,24^. This is critical for reliable temperature sensing in complex biological environments^3^. Spectral analysis revealed four peaks (P1-P4) and one valley (V1) as potential temperature calibration candidates (Fig. S3b, c). For each excitation wavelength, all 10 possible peak and valley combinations were examined. The linearity of the calibration curves was evaluated using two methods: (1) LIR calculation at specific wavelengths (one-point analysis) and (2) LIR calculation of integrated intensities across various wavelength intervals (1, 2, 3, 4, 5, and 10 nm). The obtained coefficients of determination (R^2^) are listed in Table S2. At 785 nm excitation, LIR P4/V1 (integrated intensity over 5 nm spectral range) provided the most accurate calibration, with the highest R^2^ of 0.99 for the whole thermal range and 0.97 across the biologically relevant thermal range (Fig. 3b and 3c, respectively). At 808 nm excitation, P1/P2 (integrated across 2 nm) yielded the best calibration curve, with an R^2^=0.99, and R^2^=0.97 within the lower thermal range (Fig. 3e, f), indicating a high accuracy for NP-HT applications. We attribute this high correlation to the temperature-dependent population of thermally coupled stark levels of the ^4^F_3/2_ multiplet^48^. The thermal sensitivity (SR) at 785 nm laser excitation was determined to be 0.09%/°C for the whole thermal range, and 0.15%/°C between 35-60°C. For 808 nm, SR was 0.14%/°C and 0.17%/°C for the whole range and the therapeutically relevant window, respectively. The observed results show that Nd-NC@shell particles display high sensitivity to temperature changes in the BW-I, and this observation is in agreement with previously reported studies using Nd-doped NaYF_4_ NCs, which show a similar spectral shape between 850 and 920 nm and a temperature dependent LIR trend characteristic of Nd^3+^ in a NaYF_4_ matrix^42^. Hence the as-synthesized NCs demonstrate clear potential to be used as nanothermometers for NP-HT applications.

### 3.3. Biocompatible Nd-NCs

The Nd-NCs were synthesized in an organic phase, and exhibit a surface coated with oleic acid (OA), which reduces stability in aqueous solutions and biocompatibility. For biological applications, it is essential to modify the NP surface to mitigate these effects. The addition of a silica (SiO_2)_ layer is a common approach used in biological studies because silica is biocompatible, has excellent chemical stability and is low cost. In addition, it facilitates the subsequent functionalization with different functional groups for improved uptake^49,50^. In this work, a SiO_2_ layer was added to the optimized Nd-NC@shell NCs using the reverse microemulsion technique. In this process, IGEPAL® CA-630 is used as an amphiphilic surfactant for micelle formation in an organic solvent, with hydrophobic groups on the outside and hydrophilic groups inside the micelle. The addition of OA-stabilized NP leads to a ligand exchange between OA and IGEPAL® CA-630, and the micelle is subsequently expanded upon ammonia addition to the solution. Lastly, TEOS is added to the solution and hydrolyzed, replacing IGEPAL® CA-630 and chemically absorbing on the NP surface. After condensation of the hydrolyzed TEOS, the SiO_2_ shell is formed, and the NPs can be washed and redispersed in an aqueous solution^49^.

The success of the SiO_2_-coating protocol was assessed via TEM (Fig 4a). Nd-NC@shell@SiO_2_ showed a homogeneous shape and an average size of (68 ± 8) nm. It was also possible to observe the encapsulation of single crystals inside the silica shell (shell thickness of ∼15 nm), and the absence of free SiO_2_ NPs, indicating that the TEOS:Nd-NCs ratio was adequate to coat the NC surfaces. Nd-NC@shell@SiO_2_ displayed a negative surface potential of (-46.1 ± 1.2) mV, consistent with the existence of partially deprotonated hydroxyl groups on the SiO_2_ surface^50^. To investigate the influence of SiO_2_ on emission, PL spectra of the uncoated Nd-NC@shell and coated Nd-NC@shell@SiO_2_ were compared at 785 and 808 nm excitation (0.15 mW). Figure 7 c and e show the normalized spectra ([0,1]) of Nd-NC@shell before and after SiO_2_ coating, showing no changes to the spectral shape. Hence, the previously obtained calibration curves for Nd-NC@shell can be used for thermal analysis using Nd-NC@shell@mSi NPs. Non-normalized PL spectra measured for a similar NP concentration (∼ 60 mg/mL) suggest that the addition of a SiO_2_ leads to a PL increase for both laser excitation wavelengths. Similar to what was observed for the addition of an inert NaYF_4_ shell to the active Nd^3+^-doped core, we hypothesize that the deposition of an additional protective layer leads to a passivation effect that further shields the core from solvent-induced quenching^44,47^.

**Figure 4.**
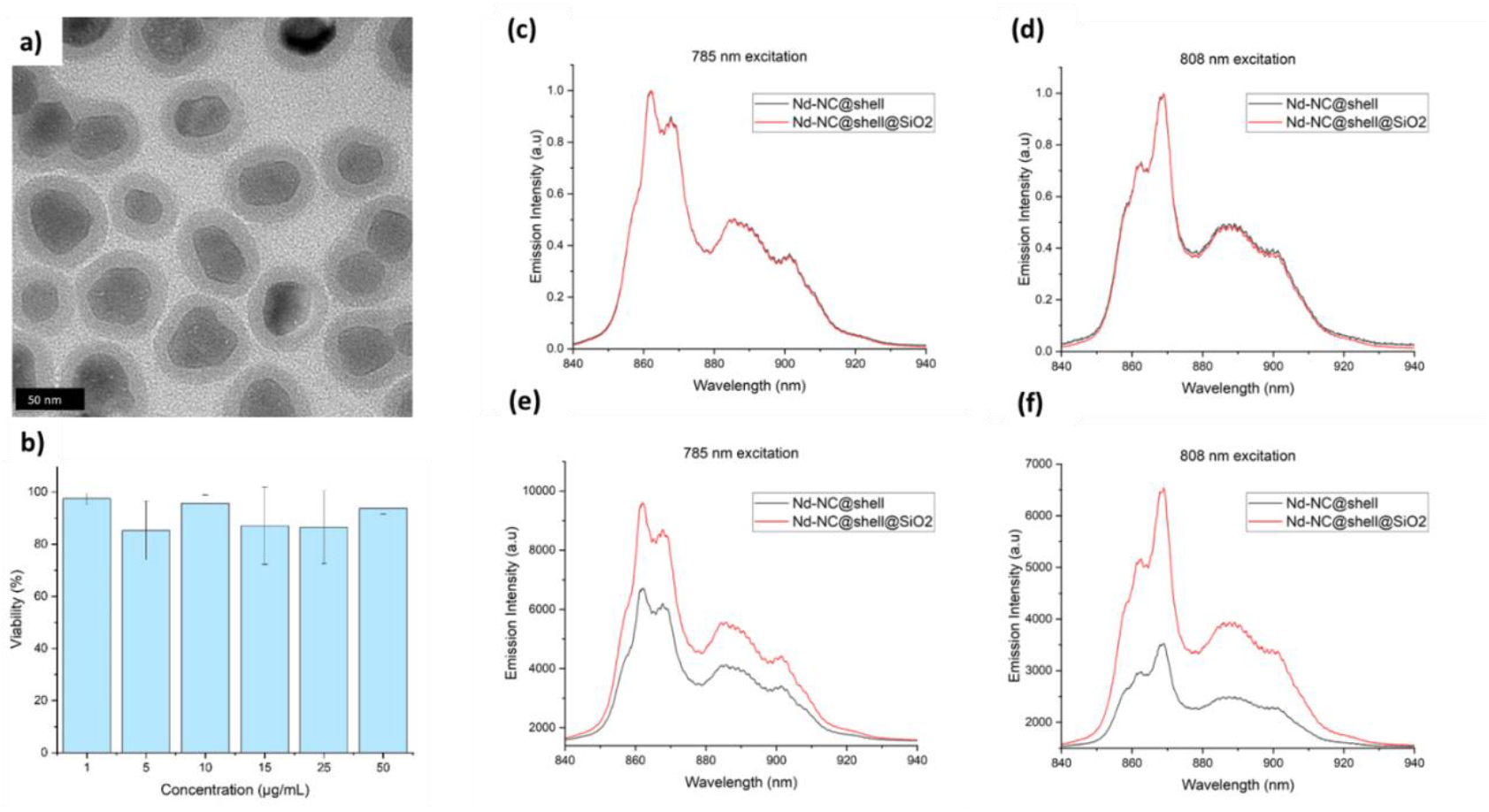
(a) TEM image of 5%Nd-NC@shell@SiO_2_. Scale bar = 50 nm; (b) Normalized PL spectra of bare and SiO_2_-coated NPs at (c) 785 and (d) 808 nm excitation; Black lines correspond to the normalized emission spectra of bare Nd-NC@shell, while Nd-NC@shell@SiO_2_ emission spectra are shown in red; Emission spectra comparing NP PL before and after SiO_2_ coating at (e) 785 and (f) 808 nm excitation.

Lastly, Nd-NC@shell@SiO_2_ toxicity was evaluated using a well-established WST-8 assay. Different concentrations of Nd-NC@shell@SiO_2_ (ranging between 1 and 50 µg/mL) were incubated with A549 lung cancer for 6 h. Figure 8 shows the calculated viability normalized for the negative control. For the tested concentrations, there was no significant decline in cell proliferation after incubation with NPs, and cell survival rate was higher than 85% for all the tested concentrations. These results are consistent with other reported studies using SiO_2_-coated NaYF_4_ NCs for similar NP concentrations^51,52^. These *in vitro* results confirm that the SiO_2_ layer endows the nanothermometers with relatively low cytotoxicity, allowing for safe thermal monitoring in biological environments.

### 3.4. Single Nanoparticle Optical Thermometry

One of the current bottlenecks of Ln-NC applications is the lack of literature data reporting the emission behavior on a single particle level. Most studies are based on the analysis of bulk samples, neglecting discrepancies in the optical properties of individual NPs^53^. Furthermore, single NP spectra enable the assessment of the sensitivity of luminescent NPs in a biologically relevant manner. Upon cellular uptake, NPs are dispersed inside the cytoplasm at low concentrations. This could potentially limit nanothermometric potential due to insufficient emission signal. To further validate the potential of the designed Nd-NCs, we performed single NP optical thermometry in NCs internalized by cancer cells. A549 cells were incubated with Nd-NC@shell@SiO_2_ (50 µg/mL) for 3 hours, and spectral mapping was performed to locate the NPs in the cells. NPs were excited using an 808 nm laser excitation. The spectral map was then compared with transmission light images of the cells, showing the overlap between the two, confirming that the NPs were located in the cell (Fig. 5a, b). Figure 5b also revealed a high internalization rate in A549 cancer cells, crucial for the application of these particles as temperature probes in NP-HT.

**Figure 5.**
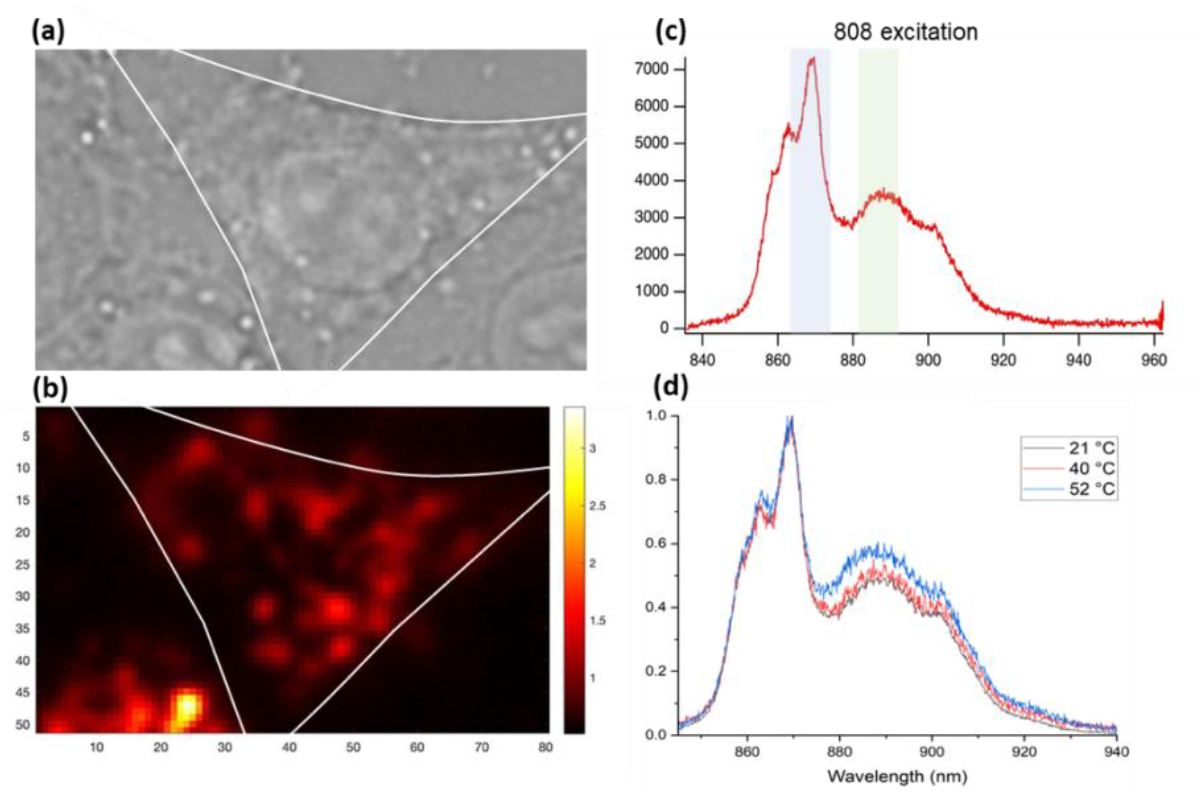
(a) Optical transmission microscopy image of A549 cells after incubation with Nd-NC@shell@SiO_2_ NPs. (b) Spectral mapping was obtained for Nd-NC@shel@SiO_2_ NPs (Ex: 808 nm. Em: 835-965 nm), showing overlap with the transmission channel. (c) Spectra was collected for single NPs, and 4 distinct peaks (P1-P4) could be detected. Spectral analysis was performed by integrating the area below the graph for P2 (868-878 nm) and P3 (885-895 nm), shown in blue and green, respectively. (d) Spectra collected at 3 distinct temperatures, showing that thermal readouts could be achieved *in vitro* in A549 cells.

Using optical spectroscopy, the emission spectra of discrete points within the cell were then collected, revealing a well-resolved spectra with clear peaks and the same spectral outline as the bulk samples. A temperature-controlled stage was then used to collect the PL spectra at different temperatures within a biological relevant range (Fig 5d). From these results, it was possible to observe a change in spectral shape with the temperature increase, indicating the potential of the Nd-NCs to detect thermal changes in biological environments for biological applications.

## 4. Conclusion and Future Prospects

In this study, we present the synthesis and characterization of biocompatible core-shell NaYF_4_ nanocrystals doped with Nd^3+^ for deep tissue nanothermometry applications, with excitation and emission wavelengths located within the first biological transparency window. Nanocrystals were prepared via thermal decomposition, resulting in a crystalline structure and homogeneous size distribution. Photoluminescence was optimized by tuning the active dopant concentration, showing that PL increases with Nd^3+^ concentration up to a certain threshold (5%). Emission was further enhanced by the addition of an inert NaYF_4_ shell. Temperature-dependence was studied by determining the luminescent intensity ratios (LIR) at different temperatures. High linear correlation was obtained, displaying the excellent thermal sensitivity of Ns-NCs at 785 and 808 nm excitation. To render the Nd-NCs biocompatible, the as-synthesized core-shell structures were coated with a silica layer, resulting in cell viabilities > 85% when incubated with up to 50 μg/ml Nd-NC@shell@SiO_2_ over 6 hours. Finally, luminescent thermal readout was demonstrated *in vitro* in A549 cells by spectrally resolving the diffraction-limited luminescence spots from single particles over a clinically relevant temperature range from 20-50 °C. The demonstration of biocompatible, nano-localized, NIR thermometry in cells is a significant step in the development of viable hyperthermal cancer treatments.

## 5. Acknowledgements

The authors are thankful to the Gobal PhD partnership program between KU Leuven and Melbourne university (GPUM/21/025). MB, SZ, PM, and JAH thank the Australian Government for funding through the Australian Research Council (ARC) Centre of Excellence in Exciton Science (CE170100026). JAH acknowledges an ARC Future Fellowship award (FT180100295). JH, HU and SR acknowledged the financial support from Research Foundation of Flanders (FWO) research grants (G0D4519N, G081916N, VS08523N), postdoctoral fellowship (for BF,12X1423N), and from the KU Leuven (C14/15/053, C14/19/079). We acknowledge the Ian Holmes Imaging Centre, Bio21 Institute, University of Melbourne for electron microscopy undertaken in this work.

