## Supporting Information for "Neodymium-Doped Nanocrystals for Diffraction-limited *in vitro* Temperature Sensing"

### Supplementary Section

Table S1| Nd-NC composition – compilation of mass of each precursor added in the synthesis of each formulation, as well as the volume of 1-octadecene (ODE) and oleic acid (OA).

| | m $_{Nd(CF3COO)3}$ (mg) | m $_{Y(CF3COO)3}$ (mg) | m $_{Na(CF3COO)3}$ (mg) | V $_{ODE}$ (mL) | V $_{OA}$ (mL) |
|---|---|---|---|---|---|
| **Neg. Control** | 0 | 538.7 | 151.2 | 10 | 10 |
| **$NaYF_4$:1%Nd** | 6.1 | 533.3 | 151.2 | 10 | 10 |
| **$NaYF_4$:2%Nd** | 12.2 | 527.9 | 151.2 | 10 | 10 |
| **$NaYF_4$:5%Nd** | 30.4 | 511.8 | 151.2 | 10 | 10 |
| **$NaYF_4$:10%Nd** | 60.8 | 484.8 | 151.2 | 10 | 10 |
| **$NaYF_4$ shell** | - | 431.0 | 137.0 | 5 | 5 |

To ensure accurate sample comparison, the laser power was kept constant (0.15 mW). To account for varying PL intensities between samples, acquisition times were adjusted. PL measurements were subsequently standardized using the calibration curves presented in Figures S2a and S2b.

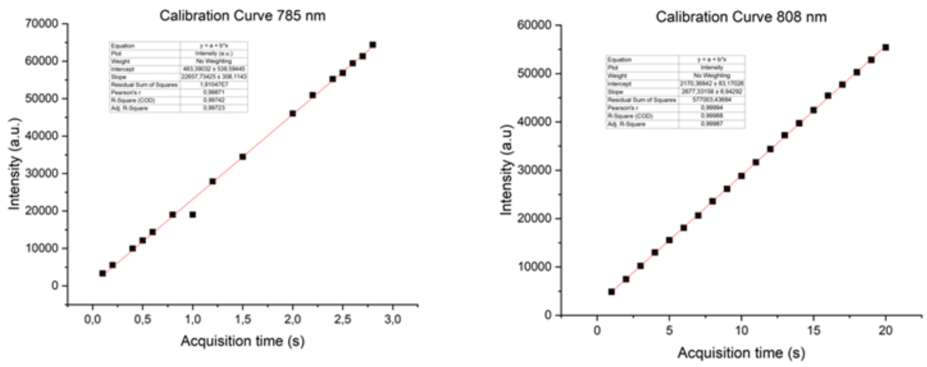


Figure S1| Calibration curves for Intensity vs . Acquisition time for (a) 785 and (b) 808 nm excitation

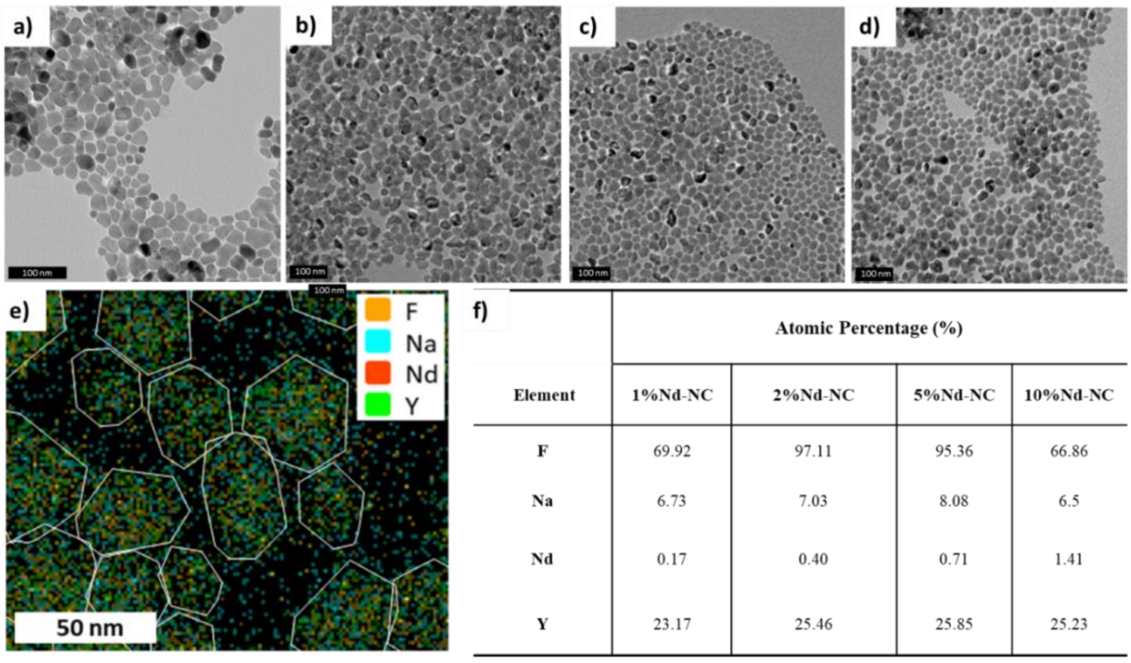


| Element | Atomic Percentage (%) 1%Nd-NC | 2%Nd-NC | 5%Nd-NC | 10%Nd-NC |
|---|---|---|---|---|
| F | 69.92 | 97.11 | 95.36 | 66.86 |
| Na | 6.73 | 7.03 | 8.08 | 6.5 |
| Nd | 0.17 | 0.40 | 0.71 | 1.41 |
| Y | 23.17 | 25.46 | 25.85 | 25.23 |

Figure S2| Transmission Electron Microscopy (TEM) images of (a) 0%Nd-NCs (b) 1%Nd-NCs (c) 2%Nd-NCs, and (d) 10%Nd-NCs. Scale bar = 100 nm; (e) EDS map of Nd-NCs showing the distribution of Nd3+ atoms inside the NPs (5%Nd-NCs@shell). NP edges were added based on SEM images of the sample. Scale bar = 50 nm; (f) Atomic Percentages of x%Nd-NC obtained from Energy Dispersive Spectroscopy (EDS).

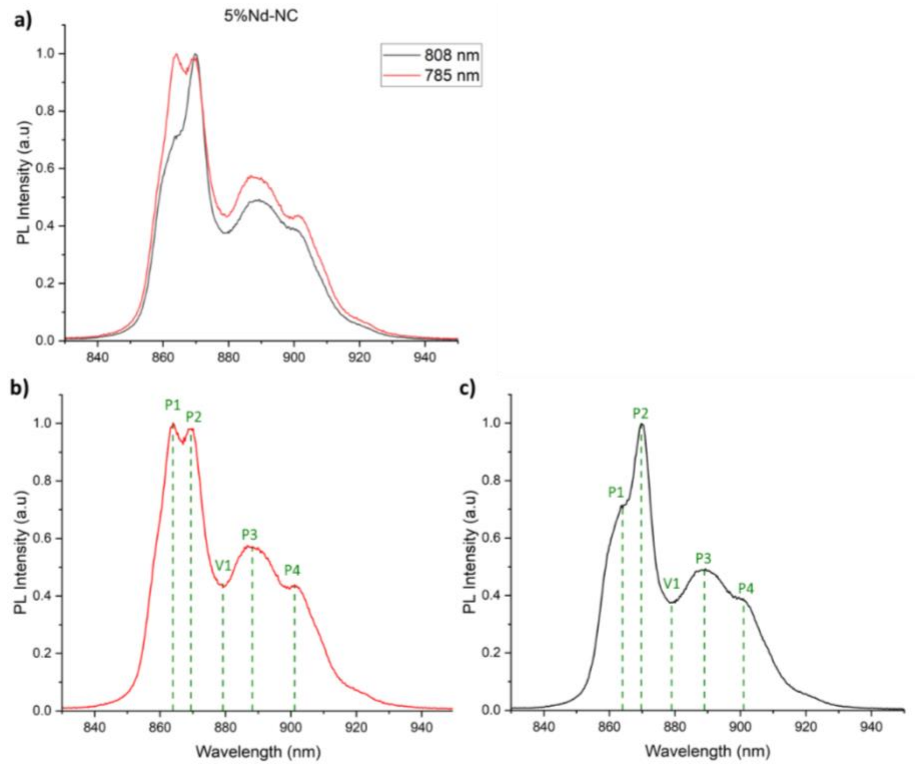


Figure S3| (a) Typical PL spectra of Nd-NCs (5%Nd-NCs) upon 785 nm and 808 nm excitation, showing spectral shape changes with the excitation wavelength. PL spectrum of 5%Nd-NCs, showing the different peaks and valleys after (b) 785 nm and (c) 808 nm excitation. Both spectra show 4 distinct peaks (P1-4) and one valley (V1), which can be used to study temperature dependence.

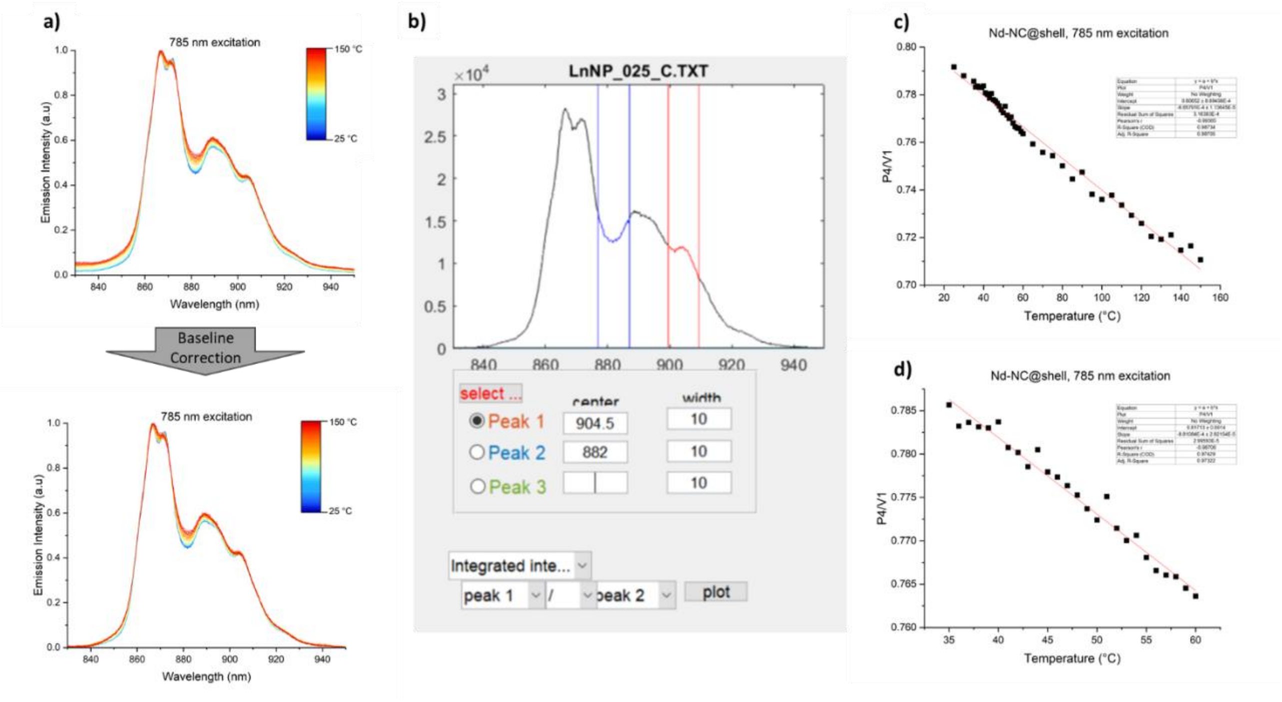


Figure S4| Temperature-dependent calibration analysis of Nd-NC@shell nanothermometers. (a) Baselined and normalized [0,1] emission spectra color-coded by temperature. (b) Matlab Graphical User Interface (GUI) used to calculate luminescence intensity ratios (LIRs) between different peaks and valleys, demonstrating the ability to adjust peak wavelengths and integration widths for analysis; (c) Full-range (25-150°C) calibration curves plotting LIR against temperature. (d) Biologically relevant (35-60°C) calibration curves.

Table S2| Coefficient of determination ($R^2$) for calibration curves obtained from various peaks (P) and valleys (V) combinations. Linearity was evaluated by calculating the intensity ratio at a specific wavelength (0 nm) and for the integrated intensity within different wavelength intervals (1, 2, 3, 4, 5, and 10 nm, unless limited by the distance between points). $R^2$ values between 0.90 and 0.95 are marked as yellow (moderate linearity), $0.95 \leq R^2 < 0.98$ is shown in light green (good linearity), and $R^2 \geq 0.98$ is shown in darker green (excellent linearity).

| **Linear Correlation (Coefficient of Determination - $R^2$)** | | | | | | | | | | | | | | |
|---|---|---|---|---|---|---|---|---|---|---|---|---|---|---|
| **Ratio** | **785 nm** | | | | | | | **808 nm** | | | | | | |
| | **0 nm** | **1 nm** | **2 nm** | **3 nm** | **4 nm** | **5 nm** | **10 nm** | **0 nm** | **1 nm** | **2 nm** | **3 nm** | **4 nm** | **5 nm** | **10 nm** |
| **P1/P2** | 0.36 | 0.60 | 0.66 | 0.62 | 0.58 | - | - | 0.96 | 0.99 | 0.99 | 0.99 | 0.99 | - | - |
| **P1/P3** | 0.81 | 0.86 | 0.85 | 0.85 | 0.85 | 0.85 | 0.86 | 0.78 | 0.90 | 0.93 | 0.92 | 0.92 | 0.91 | 0.07 |
| **P1/P4** | 0.29 | 0.33 | 0.34 | 0.33 | 0.26 | 0.24 | 0.36 | 0.92 | 0.97 | 0.97 | 0.97 | 0.97 | 0.97 | 0.93 |
| **P1/V1** | 0.94 | 0.94 | 0.94 | 0.94 | 0.94 | 0.94 | 0.94 | 0.02 | 0.30 | 0.53 | 0.56 | 0.69 | 0.83 | 0.97 |
| **P2/P3** | 0.84 | 0.90 | 0.89 | 0.89 | 0.88 | 0.87 | 0.81 | 0.86 | 0.97 | 0.98 | 0.98 | 0.98 | 0.98 | 0.96 |
| **P2/P4** | 0.54 | 0.67 | 0.65 | 0.61 | 0.53 | 0.49 | 0.16 | 0.44 | 0.85 | 0.84 | 0.86 | 0.82 | 0.69 | 0.79 |
| **P2/V1** | 0.94 | 0.95 | 0.94 | 0.94 | 0.94 | 0.94 | 0.93 | 0.96 | 0.99 | 0.99 | 0.99 | 0.97 | 1.00 | - |
| **P3/P4** | 0.67 | 0.91 | 0.91 | 0.93 | 0.95 | 0.96 | 0.96 | 0.66 | 0.91 | 0.95 | 0.96 | 0.96 | 0.96 | 0.97 |
| **P3/V1** | 0.91 | 0.96 | 0.97 | 0.97 | 0.98 | 0.98 | - | 0.79 | 0.91 | 0.95 | 0.96 | 0.96 | 0.98 | - |
| **P4/V1** | 0.96 | 0.98 | 0.98 | 0.98 | 0.98 | 0.99 | 0.99 | 0.89 | 0.97 | 0.98 | 0.99 | 0.99 | 0.99 | 0.99 |

Table S4| Coefficient of determination ($R^2$) for calibration curves obtained from various peaks (P) and valleys (V) combinations for the biological and therapeutically relevant temperature range. Linearity was evaluated by calculating the intensity ratio at a specific wavelength (0 nm) and for the integrated intensity within different wavelength intervals (1, 2, 3, 4, 5, and 10 nm, unless limited by the distance between points). $R^2$ values between 0.90 and 0.95 are marked as yellow (moderate linearity), and $0.95 \leq R^2 < 0.98$ is shown in light green (good linearity).

| **Linear Correlation (Coefficient of Determination - $R^2$)** | | | | | | | | | | | | | | |
|---|---|---|---|---|---|---|---|---|---|---|---|---|---|---|
| **Ratio** | **785 nm** | | | | | | | **808 nm** | | | | | | |
| | **0 nm** | **1 nm** | **2 nm** | **3 nm** | **4 nm** | **5 nm** | **10 nm** | **0 nm** | **1 nm** | **2 nm** | **3 nm** | **4 nm** | **5 nm** | **10 nm** |
| **P1/P2** | - | - | - | - | - | - | - | 0.68 | 0.92 | 0.97 | 0.96 | 0.96 | - | - |
| **P2/P3** | - | - | - | - | - | - | - | - | - | 0.81 | 0.80 | 0.83 | 0.80 | - |
| **P2/V1** | - | - | - | - | - | - | - | - | 0.82 | 0.85 | 0.87 | 0.92 | 0.94 | - |
| **P3/V1** | - | - | - | - | 0.85 | 0.86 | - | - | - | - | - | - | 0.61 | - |
| **P4/V1** | - | 0.95 | 0.95 | 0.95 | 0.97 | 0.97 | 0.97 | - | - | 0.71 | 0.74 | 0.82 | 0.88 | 0.91 |
